# Theoretical Design of Paradoxical Signaling-Based Synthetic Population Control Circuit in *E. coli*

**DOI:** 10.1101/2020.01.27.921734

**Authors:** Michaëlle N. Mayalu, Richard M. Murray

## Abstract

We have developed a mathematical framework to analyze the cooperative control of cell population homeostasis via paradoxical signaling in synthetic contexts. Paradoxical signaling functions through quorum sensing (where cells produce and release a chemical signal as a function of cell density). Precisely, the same quorum sensing signal provides both positive (proliferation) and negative (death) feedback in different signal concentration regimes. As a consequence, the relationship between intercellular quorum sensing signal concentration and net growth rate (cell proliferation minus death rates) can be non-monotonic. This relationship is a condition for robustness to certain cell mutational overgrowths and allows for increased stability in the presence of environmental perturbations. Here, we explore stability and robustness of a conceptualized synthetic circuit. Furthermore, we asses possible design principles that could exist among a subset of paradoxical circuit implementations. This analysis sparks the development a bio-molecular control theory to identify ideal underlying characteristics for paradoxical signaling control systems.

## Introduction

An important area in synthetic biology is the development of cell-cell communication modules that utilize coordinated feedback control to regulate population size of cell communities. Coordinated feedback control for regulation of population homeostasis has implications in engineered wound healing [1], cancer [2], and probiotic [3, 4] therapies. For the aforementioned therapies, synthetic biologists can develop genetically engineered cell systems, termed biological control circuits, that can robustly control their proliferation and/or death. Using synthetic population control, therapeutic agents can make decisions based on intercommunication between adjacent cells and the environment. Specifically for bacteria, this enhanced “sense and-respond” cell regulation capability has implications in gastrointestinal diseases caused by loss or disruption of healthy bacteria in the gut [3]. In order to alleviate this issue, one could introduce an engineered version of the disrupted bacteria that could maintain a constant cell population in the midst of environmental perturbations. Hence, research surrounding synthetic bacterial population control circuits could lead to treatments beyond traditional approaches, allowing for the development of engineered probiotic therapies. Furthermore, experimental implementation of synthetic population control designs in bacteria (in parallel with modeling and theory) can serve as prototypes to analyze underlying design principles found in engineered mammalian systems and nature.

Previous works in synthetic population control have interfaced the quorum sensing molecule N-Acyl homoserine lactone (AHL) with programmed cell death genes in *E. coli* to control the density of a cell population [5]. Within these works, experimental results show that if the “killer gene” is disconnected from the quorum sensing molecule, the population grows to carrying capacity. However, if the “killer gene” is interfaced with the quorum sensing molecule the population can grow to a size (below carrying capacity) where proliferation and death rates equal and the system reaches steady state.

In control theoretic terms, the quorum sensing molecule acts as a feedback input to suppress the growth of the cell population. This negative feedback input design yields a mono-stable population dynamic system, with one stable equilibrium point. However, the proper functioning of this feedback circuit architecture is dependent on the precise activity and expression of receptors and regulatory proteins involved within the circuit, which are susceptible to mutations. When a cell with a receptor loss-of-function mutation (exhibiting a dysregulated proliferative response) arises, it may invade the cell population and thus break the homeostatic control [6]. A paradoxical feedback input design has been shown to prevent such a takeover in natural occurring mammalian systems [6–9].

In population control via paradoxical signaling, the same quorum sensing signal is interfaced with both proliferative and apoptotic cell-intrinsic mechanisms, providing both positive and negative feedback in different signal concentration regimes. With this feedback circuit architecture, cells exhibit an apoptotic or proliferative response when below or above a signal threshold value. If above the aforementioned signal threshold, cells are driven to a high steady state cell population. Conversely, if below the signal threshold, cells are driven to a low steady state cell population. Thus, this paradoxical feedback input design can yield a bi-stable cell population dynamic system, with one unstable and two stable equilibrium points [6–9]. The paradoxical feedback control circuit allows for cell intrinsic “safety mechanisms" against aberrant proliferation, due to receptor loss-of-function mutations. Furthermore, it can be shown that the paradoxical architecture provides increased stability in the presence of environmental perturbations [9]. Another benefit of the bistable property of the paradoxical feedback architecture, is the possibility to have two homeostatic states such that the cell population could expand or collapse to a specified size when needed, which could be necessary for the aforementioned applications.

Here, we develop a mathematical model representing the behavior of a synthetic paradoxical circuit design in *E. coli* through a set of nonlinear differential equations describing the known extracellular signaling and intracellular reactions that drive the circuit function. The developed model is motivated by previous mathematical models of natural mammalian paradoxical systems [6–9] and ongoing experimental implementations of synthetic mammalian and bacterial systems. Our modeling approach is distinct from previous works, which are based in heuristics, since we take a “ground up” approach by developing detailed differential equations of significant reactions and then simplifying them using singular perturbation techniques. We use our developed model to explore performance and robustness properties in a control theoretic context.

## Results

In what follows, we elaborate on the circuit components of the proposed synthetic paradoxical feedback system in *E. coli* and corresponding reactions. We also describe the corresponding differential equation model and use singular perturbation theory for model order reduction. We also explore the conditions under which the circuit components within the paradoxical signaling architecture yield bi-stable population dynamics necessary for mutational robustness. Some additional properties, such as stability, are also discussed.

### Paradoxical Circuit Components and Corresponding Reactions

We derived a biochemical model of the paradoxical signaling population control gene synthetic circuit implementation. The set of biochemical reactions are shown below in equations (1)–(6):

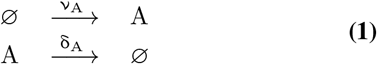

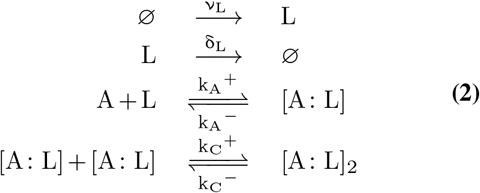

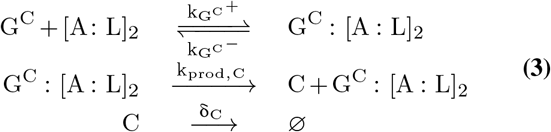

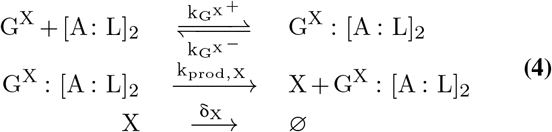

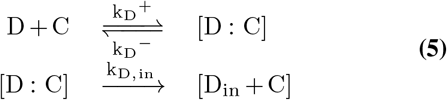

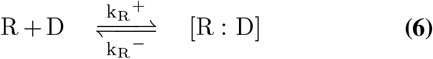

The paradoxical feedback circuit architecture (schematic description shown in Fig. 1a) consists of a Lux-AHL quorum sensing system (3OC6HSL autoinducer with cytoplasmic receptor, LuxR) interfaced with both lysis protein (X174) gene and drug resistance (kanamycin resistance) gene. As shown, the net effect of AHL induced expression of X174 (which promotes cell lysis) is to provide negative feedback that suppresses cell proliferative response. Conversely, AHL induced kanamycin resistance provides positive feedback since increased resistance promotes cell proliferative response in the presence of kanamycin drug. Following synthesis with a constitutive promoter, 3OC6HSL (AHL) diffuses freely in and out of the cell and its concentration increases as the cell density of the population increases. Reactions in equation set (1) describe AHL (denoted by *A*) production and degradation. AHL binds to its LuxR cytomasimic receptor (denoted by *L*), and forms the LuxR-AHL complex (denoted by [A:L]) which dimerizes and then activates PLux promoter present on both X174 gene and kanamycin resistance gene (denoted by *G*^*X*^ and *G*^*C*^ respectively). Without the AHL ligand, the LuxR protein is unstable and is rapidly degraded [10]. Reactions in equation set (2) describe the formation of the LuxR-AHL complex.

**Fig. 1.**
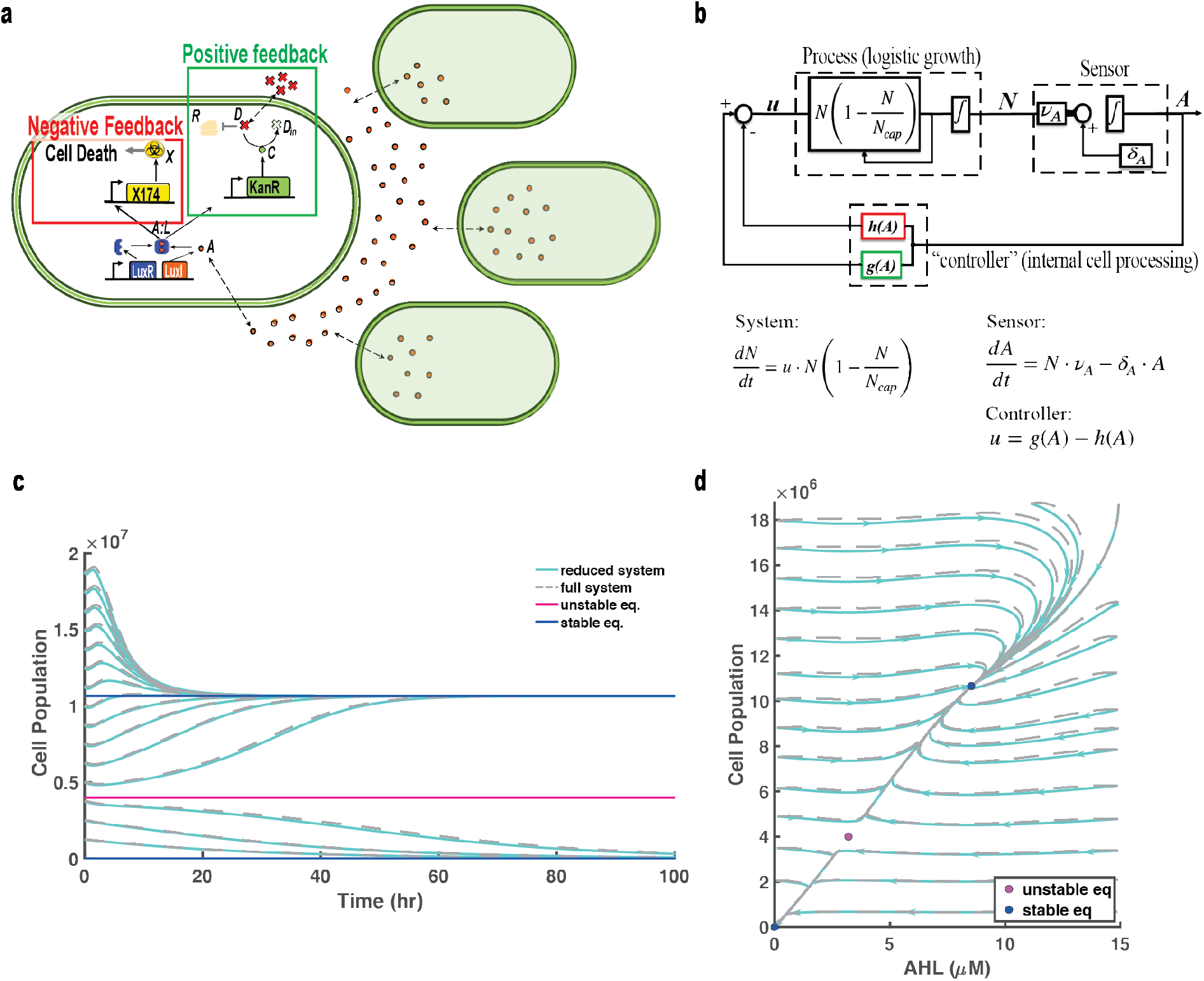
**a.** Schematic description of paradoxical feedback circuit architecture. The circuit consists of Lux-AHL quorum sensing system with 3OC6HSL autoinducer (denoted by *A*) with cytoplasmic receptor, LuxR (denoted by *L*). The quorum sensing system is interfaced with both lysis protein (X174, denoted by *X*) gene and drug resistance (kanamycin resistance, denoted by *C*) gene. **b.** The paradoxical feedback circuit acts as a feedback control system of the dynamics of the cell population. *u* is the signal dependent net growth rate which acts as a feedback control input to the dynamics of the cell population. The change in population size (process dynamics) can be sensed by the change in quorum sensing signal (sensor dynamics) produced and released by each cell. The signal is fed back to each cell for internal processing (controller). **c.** Simulation comparison of the reduced system (equations shown in b.) against the full system (shown in equations (7) - (12)). Both systems reproduce the bi-stable dynamics characteristic of paradoxical signaling. The reduced dynamics obtained from the singular perturbation approach (cyan solid line) well represent the dynamics of the full system (grey dotted line). **d.** Phase diagram of the reduced (cyan solid line) and full (grey dotted line) systems further verifies that three equilibrium points are possible with the higher and lower points being stable and middle point being unstable.

Finally, reactions in equation sets (3) and (4) describe the subsequent expression of X174 and kanamycin resistance (denoted by *X* and *C* respectively). Reactions in equation set (5) describe inactivation of kanamycin (denoted by *D*) by kanamycin resistance. Reactions in equation set (6) describes how kanamycin binds to ribosomes (denoted by *R*) which inhibits ribosomal function [11]. Species degradation is denoted as ∅.

The set of reaction equations (1)–(6) were obtained under the following assumptions:

1. AHL diffuses rapidly inside and outside the cell [5].
2. The production rate, *ν*_*A*_, of AHL is assumed to be constant (or depends on variables that do not change in time and hence are left outside the model) [12].
3. The degradation rates (*δ_A_, δ_L_, δ_C_, δ_X_*) of *A*, *L*, *C* and *X* are assumed to be constant for each individual molecule of the corresponding species [12, 13].
4. Formation of LuxR-AHL complex and subsequent dimerization are reversible reactions [12, 13].
5. Binding of kanamycin to kanamycin resistant protein is reversible [11, 14].
6. Binding of ribosome to kanamycin is reversible [15].

### Derivation of Differential Equation Model

The dynamical evolution of biochemical reaction equations (1)–(6) was obtained using the mass-action kinetics formalism [16]. Equations (7)–(11) represent the dynamics of each species inside a cell within a homogeneous cell population:

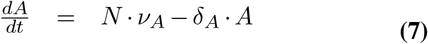

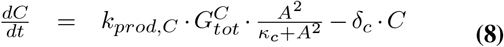

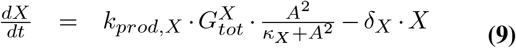

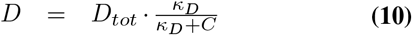

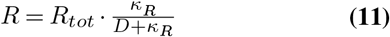

Here *κ*_*C*_, *κ*_*X*_, *κ*_*R*_, *κ*_*D*_ are constant expressions containing parameters in reaction equations (2) - (6) as explained in Supplementary Note 1. The following assumptions are used for the aforementioned equations:

1. Binding and enzymatic reactions in equation (15) are fast with respect to AHL production and degradation reactions [16, 17]. Therefore the binding reactions in equations (2)–(4) have reached steady state.
2. The total amount of kanamycin drug (*D*_*tot*_), DNA 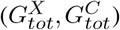, and ribosomes (*R_tot_*) is conserved.
3. The amount of inactivated kanamycin drug (*D_in_*) is small compared to total amount of drug (*D*_*tot*_).

In equation (7), we incorporate the global AHL signal by summing the contribution of AHL production from each cell. Here, *N* represents the time varying number of cells within the population. Equation (12) describes the net effect on the rate of change of cell population (*N*) assuming that the growth rate of bacteria is dependant on the number of ribosomes (unbound to kanamycin) in the cell [18] and the death rate is directly proportional to the concentration of lysis protein [5]. Parameters *ψ*_1_ and *ψ*_2_ are maximum and minimum proliferation and death rates respectively. Parameter *δ*_*N*_ is the lysis protein dependant death rate. Equation (7) and (12) represent the coupling between local detailed mechanisms, global signal, and dynamic change in population.

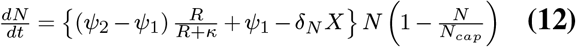

### Model Order Reduction using Singular Perturbation Theory

Using singular perturbation theory [19], we may approximate a reduced system where the dynamics of the slow variables are considered. We consider the protein characteristic degradation (of kanamycin resistance and X174 proteins) to be ~.*7hr*^−1^, which can be achieved with the use of degradation tags [20, 21]. This rate is an order of magnitude more than AHL degradation of ~.02*hr*^−1^ [22]. Let *ϵ* := *δ*_*A*_/*δ*_*X*_, which is the ratio of AHL degradation to X174 (characteristic) degradation. We may rewrite the above system by using the definitions:

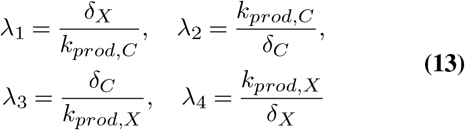

This follows that:

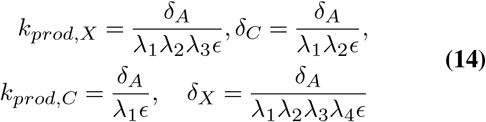

So we obtain:

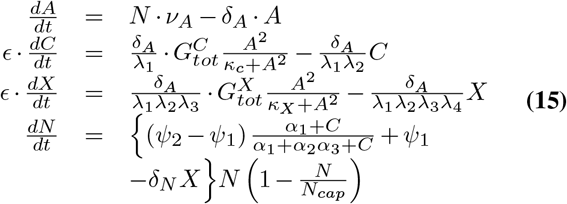

Where we have substituted equations (10) and (11) into equation (12). Constants *α*_1_, *α*_2_ and *α*_3_ are constant expressions containing parameters in reaction equations (5) and (6) as explained in Supplementary Note 1.

If AHL degradation is slow compared to the characteristic protein degradation rate, *E* can be approximated as zero (*E* ≪ 1 ⇒ *E* ≈ 0) and we may write the reduced system as:

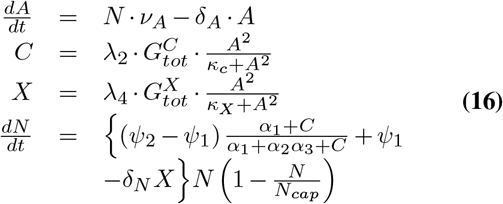

Notice the first term on the right hand side describing the rate of change of cell number 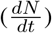 is a Hill-like function [17] that is a monotonically increasing since *α*_3_ > 1 (see Supplementary Note 1 for expression). This implies that higher kanyamycin resistance leads to increased proliferation.

Final substitutions and simplifications give:

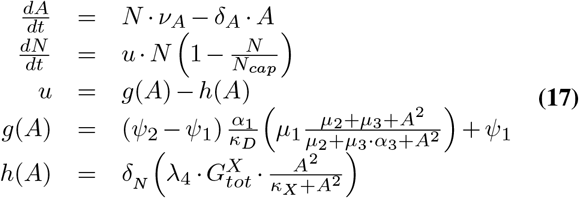

The reduced system can approximate the full system if the boundary layer system reaches asymptotic stability [16], which is verified in Supplementary Note 2. Here, *μ*_1_, *μ*_2_,*μ*_3_ are constant expressions as explained in Supplementary Note 1.

### Reduced Order Model can Reproduce Characteristic Properties of Paradoxical Feedback Population Control System

As previously mentioned, the circuit acts as a closed loop feedback control system of the dynamics of the cell population. Considering the block diagram in Fig. 1b, function *u* represents the signal dependent net growth rate *g*(*A*) − *h*(*A*) of the cell population (*N*). *u* acts as a feed-back control input to the dynamics of the cell population. The change in population size (process dynamics) can be sensed by the change in quorum sensing signal (sensor dynamics) being produced and released by each cell. The signal is fed back to each cell for internal processing (controller). The structure and parameters of functions *g*(*A*) and *h*(*A*) that comprise the single-cell level controller are dictated by intracellular reaction mechanisms.

As seen in Fig. 1c, by simulating the reduced system we are able to reproduce the bi-stable dynamics characteristic of paradoxical signaling using characteristic biological parameters. Furthermore the reduced system can accurately represent the full system. Regardless of initial population, the cell number converges to either the higher or lower steady state. We further verify that three equilibrium points are possible (phase diagram in 1d) with the middle point being unstable, and the higher and lower points being stable. (See verification of equilibrium point stability in Supplementary Note 3.)

### Model shows that Mutated Cells with Paradoxical Feedback are at a Growth Disadvantage

Fig. 2 demonstrates the resilience of the paradoxical feedback architecture to cytoplasmic receptor LuxR loss-of-function mutations. To represent the paradoxical feedback circuit, the system in equation (17) is simulated using characteristic parameters. For the mutant case, variable *A*, representing AHL, is set to zero for all time in functions *g*(*A*) and *h*(*A*). This is in order to represent that although the signal is being produced it cannot be sensed by the internal cell controller. For the non-paradoxical (negative feedback) circuit, function *g*(*A*) is set to zero to represent pure negative feedback.

**Fig. 2.**
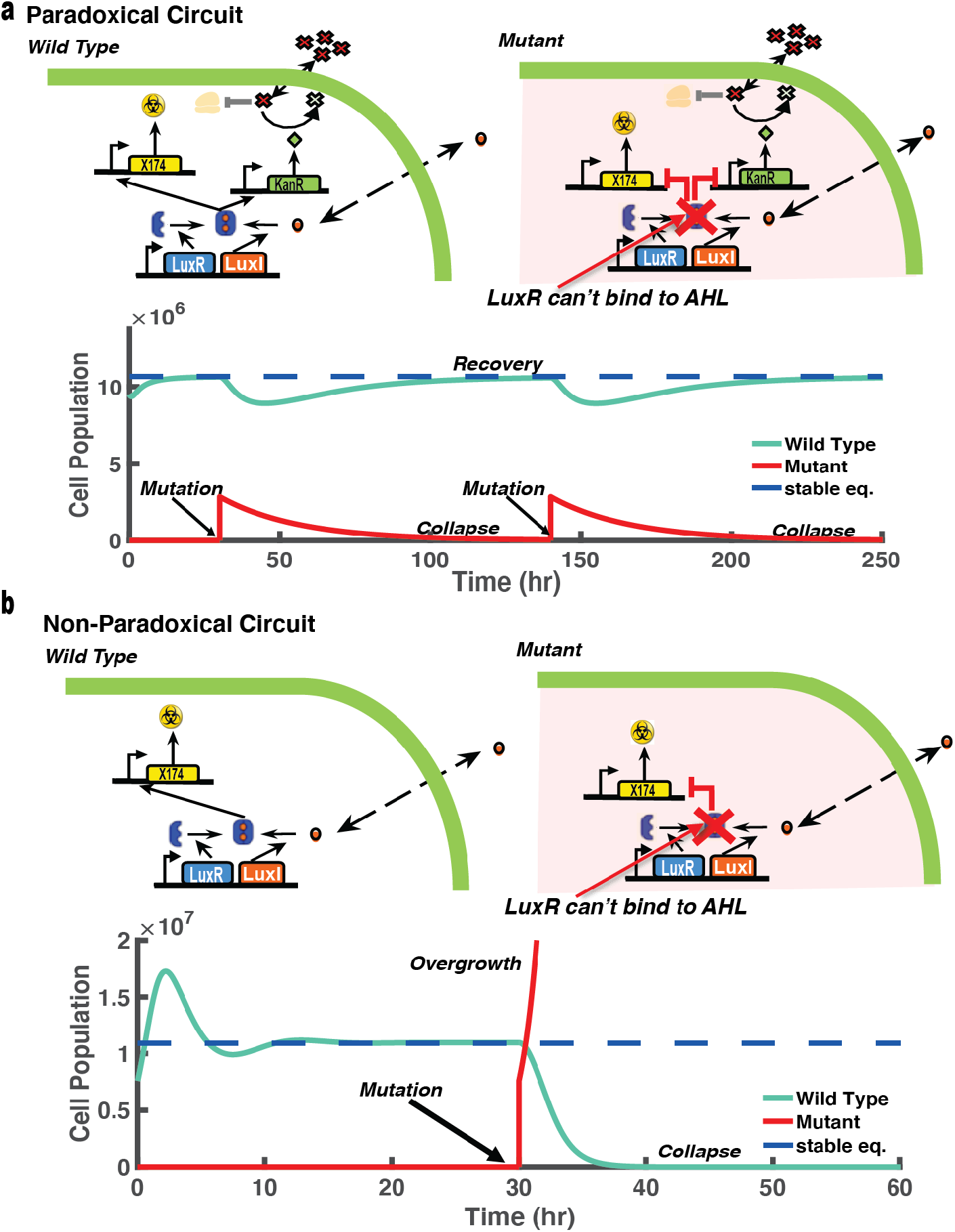
**a.** Simulation of mutated cells with paradoxical feedback show that they are at a growth disadvantage since the LuxR-AHL complex can no longer be formed, which effectively blocks expression of X174 and kanamycin resistance. This leaves mutated cells vulnerable to kanamycin induced death. **b.** Simulation of mutated cells with non-paradoxical (purely negative) feedback are at a growth advantage since the cells can no longer produce the lysis protein necessary to suppress proliferation.

As shown in Fig. 2, mutated cells with paradoxical feed-back are at a growth disadvantage since the LuxR-AHL complex can no longer be formed. This effectively blocks activation of PLux promoter and subsequent expression of X174 lysis protein and kanamycin resistance. Consequently, mutant paradoxical feedback cells are vulnerable to kanamycin induced death. We also consider the case of cells with non-paradoxical (purely negative) feedback, which controls growth through AHL induced lysis protein expression in the absence of drug and drug resistance gene [5]. Mutated cells of this type are at a growth advantage since the cells no longer produce the lysis protein necessary to suppress proliferation.

### Circuit Components Effect Bi-Stability Properties of Paradoxical Feedback System

By understanding how the controller is affected by paradoxical signaling implementations it is possible to improve system performance and robustness properties and meet pre-specified design objectives. Previous mathematical analysis of natural paradoxical feed-back mechanisms in T-cells suggested possible design principles of the paradoxical architecture [9]. Namely, altering intracellular mechanisms to increase positive feedback within the paradoxical circuit may lead to increased stability in the population dynamic system. This is because increased positive feedback increases the basin of attraction of the highest stable equilibrium point. Furthermore, intracellular mechanisms that exhibit ultra-sensitive (high-order non-linear) signal detection sensitivity are necessary for the desired bi-stable population dynamic response (i.e two stable, highest and lowest, equilibrium points with one, middle, unstable equilibrium point).

In order to verify the effect of increased positive feed-back on the current synthetic paradoxical architecture we varied the total amount of kanamycin (*D*_*tot*_) within the system, which effectively varied the rate of proliferative response and thus positive feedback. As seen in Fig. 3a, lowering the amount of kanamycin (effectively increasing positive feed-back) causes the middle and highest equilibrium point of the population dynamic system to shift away from each other effectively increasing the basin of attraction at the highest equilibrium point.

**Fig. 3.**
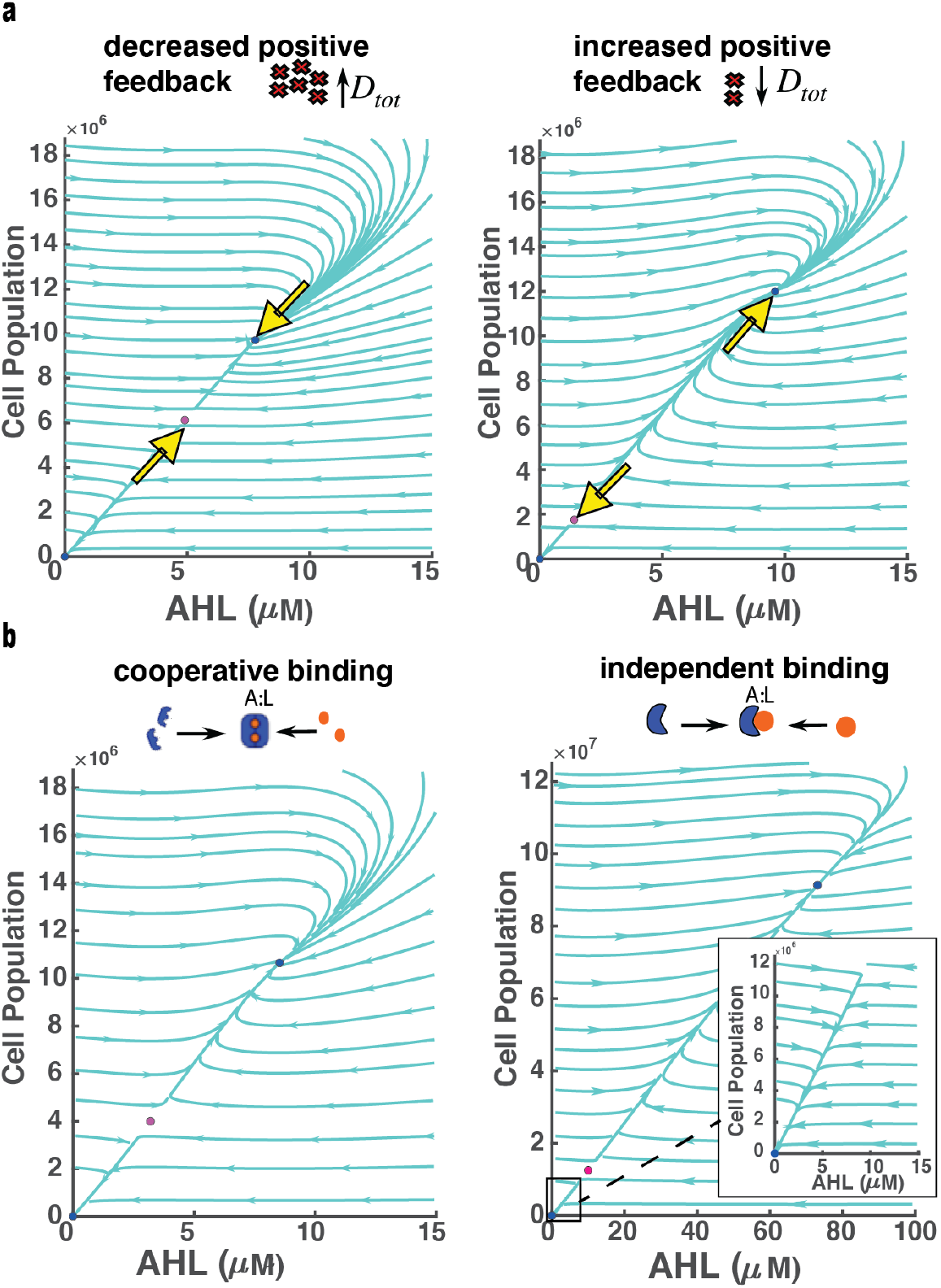
**a.** (left) Increasing the amount of kanamycin (i.e. effectively decreasing positive feedback) causes the middle and highest equilibrium point to shift towards each other. (right) Lowering the amount of kanamycin (i.e. effectively increasing positive feedback) causes the middle and highest equilibrium point to shift away from each other effectively increasing the basin of attraction of the highest equilibrium point. **b.** Simulation of internal cell (controller) mechanisms with independent binding of LuxR-AHL complex (right) show order-of-magnitude increase in middle and high equilibrium points. However, when considered within the AHL regime of the original cooperative binding circuit (left), population dynamics appear mono-stable (inset).

Positive cooperative binding (i.e. dimerization) of the LuxR-AHL complex produces higher-order nonlinearities represented by hill functions with hill coefficient greater than one, as seen in equations (8) and (9). In order to verify the effect of higher order nonlinearities on population dynamics, we consider a scenario with independent binding of LuxR-AHL complex, where dimerization of LuxR-AHL does not occur before activation of the PLux promoter. Under these conditions the hill coefficient of the associated hill function changes to one. The resulting simulations in this case, Fig. 3, show an order-of-magnitude increase in middle and high equilibrium points. Therefore, internal cell processing mechanisms represented by hill functions with hill coefficient equal to one are capable of exhibiting bi-stable population dynamic characteristics. In this case, when similar rate parameters are considered between the two systems, the bi-stability regime of the LuxR-AHL independent binding system is significantly extended. Furthermore, when the LuxR-AHL independent binding system is considered within the AHL regime of the original cooperative binding circuit the population dynamics appear mono-stable, as seen in the inset of Fig. 3.

## Discussion

In the development and implementation of a synthetic paradoxical feedback circuit, several fundamental questions arise. Specifically, what are the possible intracellular mechanisms that lead to paradoxical signaling feedback control? And how can these mechanisms be regulated to improve stability, robustness and performance of population control in therapeutic contexts? In this paper, we presented a mathematical analysis of a synthetic paradoxical feedback circuit for bacterial population control. We distinguished our analysis from existing theoretical studies of natural paradoxical feedback mechanisms in T-cells by taking a mechanistic and control theoretic approach, which elucidated possible design requirements for stability and robustness properties. The presented results are a first step in development of a bio-molecular control theory for paradoxical feedback systems. Further development of a bio-molecular control theory to study this system could elucidate efficient and systematic rational design of synthetic paradoxical feedback circuits while enabling a deeper understanding of fundamental design principles found in nature.

The proposed paradoxical feedback controller may have some draw-backs in the case of mutations in X174 expression, which would cause aberrant proliferation. However, in this case, the presence of kanamycin drug (with still functioning kanamycin resistance) would keep mutated cells from growing to carrying capacity. Even so, the proposed paradoxical feedback architecture retain advantages over previous negative feedback architectures.

This analysis can also give some insight when initially considering components for construction of a synthetic paradoxical feedback circuit in bacteria. Specifically, expressing drug resistance in *E. coli* while varying the exogenous amount of drug, may allow for tune-able positive feedback within the paradoxical circuit architecture. This could be used to vary stability properties and fine-tune population control set points. Furthermore, circuit implementations containing alternate quorum sensing systems with different cooperative binding behaviors could be considered given the desired design objectives. Possibly, quorum sensing systems with higher-order nonlinear characteristics could be used to constrain bi-stable population characteristics to within a certain regime. Ultra-sensitivity also could be incorporated by other mechanisms. For example, initiation of AHL production through activation of AHL sensitive promoter could also lead to ultra-sensitive characteristics. Consideration of these alternatives re-enforce the need for continued bio-molecular control theory development and mathematical analysis to understand how paradoxical signaling regulated proliferation/death mechanisms lead to the observed population behaviors.

## ACKNOWLEDGEMENTS

The authors are grateful to Leopold N. Green for help with conceptualizing circuit components within design and elucidating mechanisms. The authors acknowledge funding support from the Burroughs Wellcome Fund. The efforts depicted is also sponsored by the Defense Advanced Research Projects Agency (Agreement HR0011-17-2-0008). The content of the information does not necessarily reflect the position or the policy of the Government, and no official endorsement should be inferred.

## Supplementary Note 1 Summary of Constant Expressions and Parameter Values

**Table 1.** Constant Expressions.

| Constant | Expression | unit |
| --- | --- | --- |
| $\kappa_C$ | $\left(\frac{\delta_L}{\nu_L}\right)^2 \cdot \frac{k_{GC}^-}{k_{GC}^+} \cdot \frac{k_C^-}{k_C^+} \cdot \left(\frac{k_A^-}{k_A^+}\right)^2$ | $\mu M^2$ |
| $\kappa_X$ | $\left(\frac{\delta_L}{\nu_L}\right)^2 \cdot \frac{k_{GX}^-}{k_{GX}^+} \cdot \frac{k_C^-}{k_C^+} \cdot \left(\frac{k_A^-}{k_A^+}\right)^2$ | $\mu M^2$ |
| $\kappa_R$ | $\frac{k_R^-}{k_R^+}$ | $\mu M$ |
| $\kappa_D$ | $\frac{k_{D,in} + k_D^-}{k_D^+}$ | $\mu M$ |
| $\alpha_1$ | $\frac{R_{tot}}{R_{tot} + \kappa} \kappa_D$ | $\mu M$ |
| $\beta$ | $\frac{\kappa_D}{\kappa_R} \cdot (D_{tot} + \kappa_R)$ | $\mu M$ |
| $\alpha_2$ | $\frac{\kappa}{R_{tot} + \kappa} \beta$ | $\mu M$ |
| $\alpha_3$ | $\frac{D_{tot} + \kappa_R}{\kappa_R}$ | unit-less |
| $\mu_1$ | $\frac{\alpha_1 + \alpha_2 + \lambda_2 G_{tot}^C}{\alpha_1 + \alpha_2 \alpha_3 + \lambda_2 G_{tot}^C}$ | unit-less |
| $\mu_2$ | $\kappa_C \frac{(\alpha_1 + \alpha_2)(\alpha_1 + \alpha_2 \alpha_3) + (\lambda_2 G_{tot}^C)(\alpha_1)}{(\alpha_1 + \alpha_2 + \lambda_2 G_{tot}^C)(\alpha_1 + \alpha_2 \alpha_3 + \lambda_2 G_{tot}^C)}$ | $\mu M^2$ |
| $\mu_3$ | $\kappa_C \frac{(\lambda_2 G_{tot}^C)(\alpha_2)}{(\alpha_1 + \alpha_2 + \lambda_2 G_{tot}^C)(\alpha_1 + \alpha_2 \alpha_3 + \lambda_2 G_{tot}^C)}$ | $\mu M^2$ |

**Table 2.**
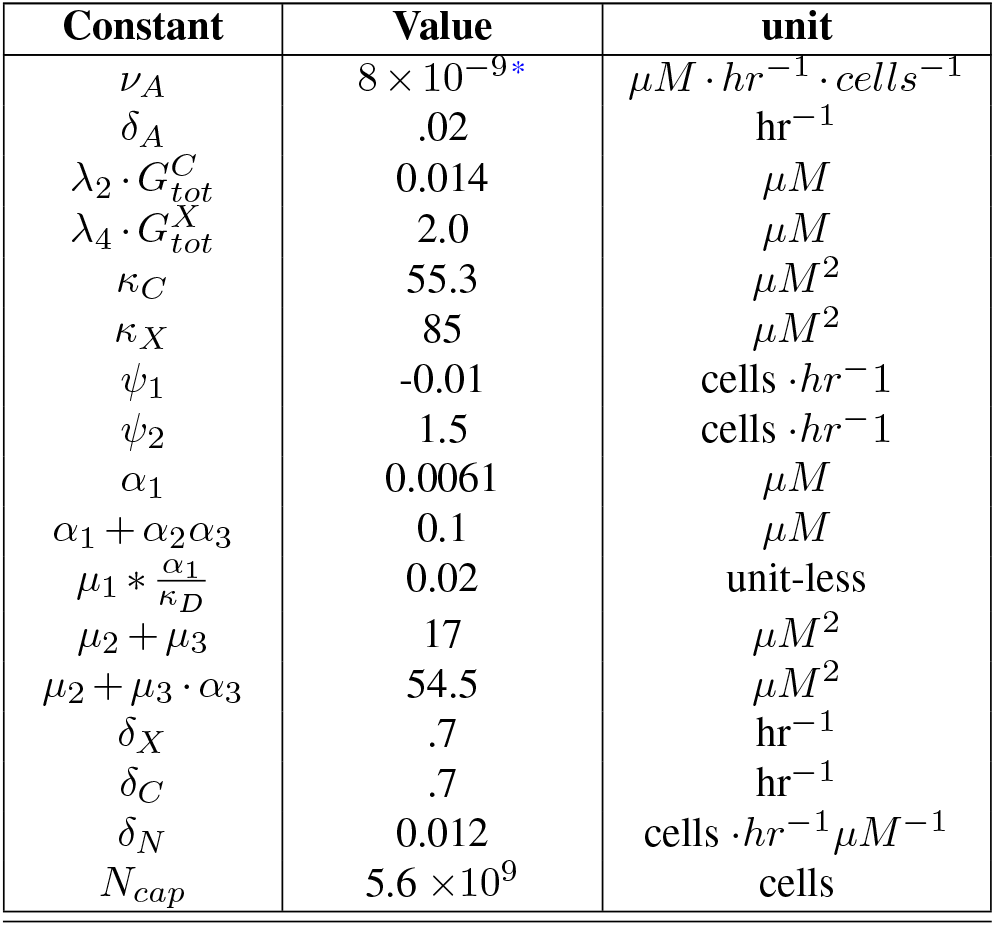
Parameter Values.

## Supplementary Note 2 Boundary Layer Analysis

Consider the boundary layer system of system in equation (17):

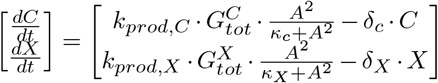

∗This value takes into consideration intracellular and extracellular AHL by incorporating the ratio between the cellular and the environment volumes [12].

The reduced system can approximate the full system if the boundary layer system reaches asymptotic stability, which is verified by calculating the eigenvalues of the linearized system evaluated at slow manifold, 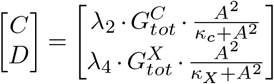. The condition for stability is [16]:

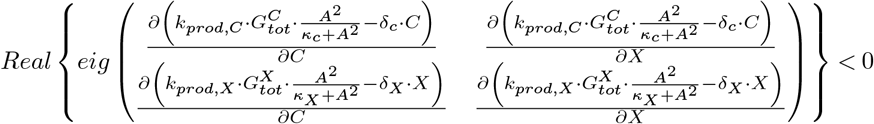

Which states that the eigenvalues of the Jacobian for the boundary layer system must have a negative real part. Solving for the eigenvalues:

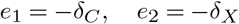

The eigenvalues are negative and real for all biologically relevant situations.

## Supplementary Note 3 Verification of Equilibrium Point Stability in Full System

We calculate the eigenvalues of the linearized systems about their equilibrium points. The Jacobian of the full system defined in equations (7) – (12) is:

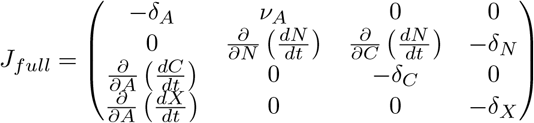

Calculating the eigenvalues for the given parameter values at the highest equilibrium point:

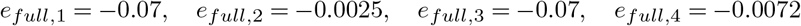

Calculating the eigenvalues for the given parameter values at the lowest equilibrium point:

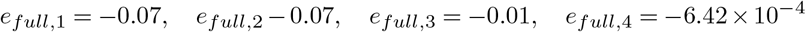

Since all the eigenvalues are negative and real, this guarantees asymptotic stability at the highest and lowest equilibrium point.

